# Latent human herpesvirus 6 is reactivated in chimeric antigen receptor T cells

**DOI:** 10.1101/2022.08.12.503683

**Authors:** Caleb A. Lareau, Yajie Yin, Katie Maurer, Katalin D. Sandor, Garima Yagnik, José Peña, Jeremy Chase Crawford, Anne M. Spanjaart, Jacob C. Gutierrez, Nicholas J. Haradhvala, Tsion Abay, Robert R. Stickels, Jeffrey M. Verboon, Vincent Liu, Jackson Southard, Ren Song, Wenjing Li, Aastha Shrestha, Laxmi Parida, Gad Getz, Marcela V. Maus, Shuqiang Li, Alison Moore, Rafael G. Amado, Aimee C. Talleur, Paul G. Thomas, Houman Dehghani, Thomas Pertel, Anshul Kundaje, Stephen Gottschalk, Theodore L. Roth, Marie J. Kersten, Catherine J. Wu, Robbie G. Majzner, Ansuman T. Satpathy

## Abstract

Cell therapies have yielded durable clinical benefits for patients with cancer, but the risks associated with the development of therapies from manipulated human cells are still being understood. For example, we currently lack a comprehensive understanding of the mechanisms of neurotoxicity observed in patients receiving T cell therapies, including recent reports of encephalitis caused by human herpesvirus 6 (HHV-6) reactivation^1^. Here, via petabase-scale viral RNA data mining, we examine the landscape of human latent viral reactivation and demonstrate that HHV-6B can become reactivated in human CD4+ T cells in standard *in vitro* cultures. Using single-cell sequencing, we identify a rare population of HHV-6 ‘super-expressors’ (~1 in 300-10,000 cells) that possess high viral transcriptional activity in chimeric antigen receptor (CAR) T cell culture before spreading to infect other cells *in vitro*. Through the analysis of single-cell sequencing data from patients receiving cell therapy products that are FDA-approved^2^ or used in clinical studies^3,4^, we identify the presence of CAR+, HHV-6 super-expressor T cells *in vivo*. Together, our study implicates cell therapy products as a source of lytic HHV-6 reported in clinical trials^1,5–7^ and has broad implications for the design, production, and monitoring of cell therapies.

## Main

During the first years of life, humans become exposed to endemic pathogens that infect nearly the entire population^8,9^. Although primary infection typically clears with sub-clinical symptoms, viruses from the *Herpesviridae, Polyomaviridae, Adenoviridae*, and *Parvoviridae* families, among others, can become ‘quiescent,’ resulting in a latent phase of the virus that is stably maintained in healthy individuals^9^. However, under acute stress conditions, such as fever, immunosuppressive states such as hematopoietic stem cell transplant (HSCT)^10^, or trauma^8^, latent viruses can become reactivated, leading to a variety of complex clinical manifestations^11^ (**Fig. 1a**). Though anecdotes of latent viral reactivation have been reported, we lack a complete understanding of the factors and mechanisms underlying latency and the settings in which they become reactivated. Notably, viral reactivation is a major clinical side effect of HSCT^10^ and more recently described as a complication in patients receiving CAR-T cell therapies^1^. However, the factors driving viral latency and reactivation in these emerging cellular therapeutic modalities have not been characterized.

**Figure 1.**
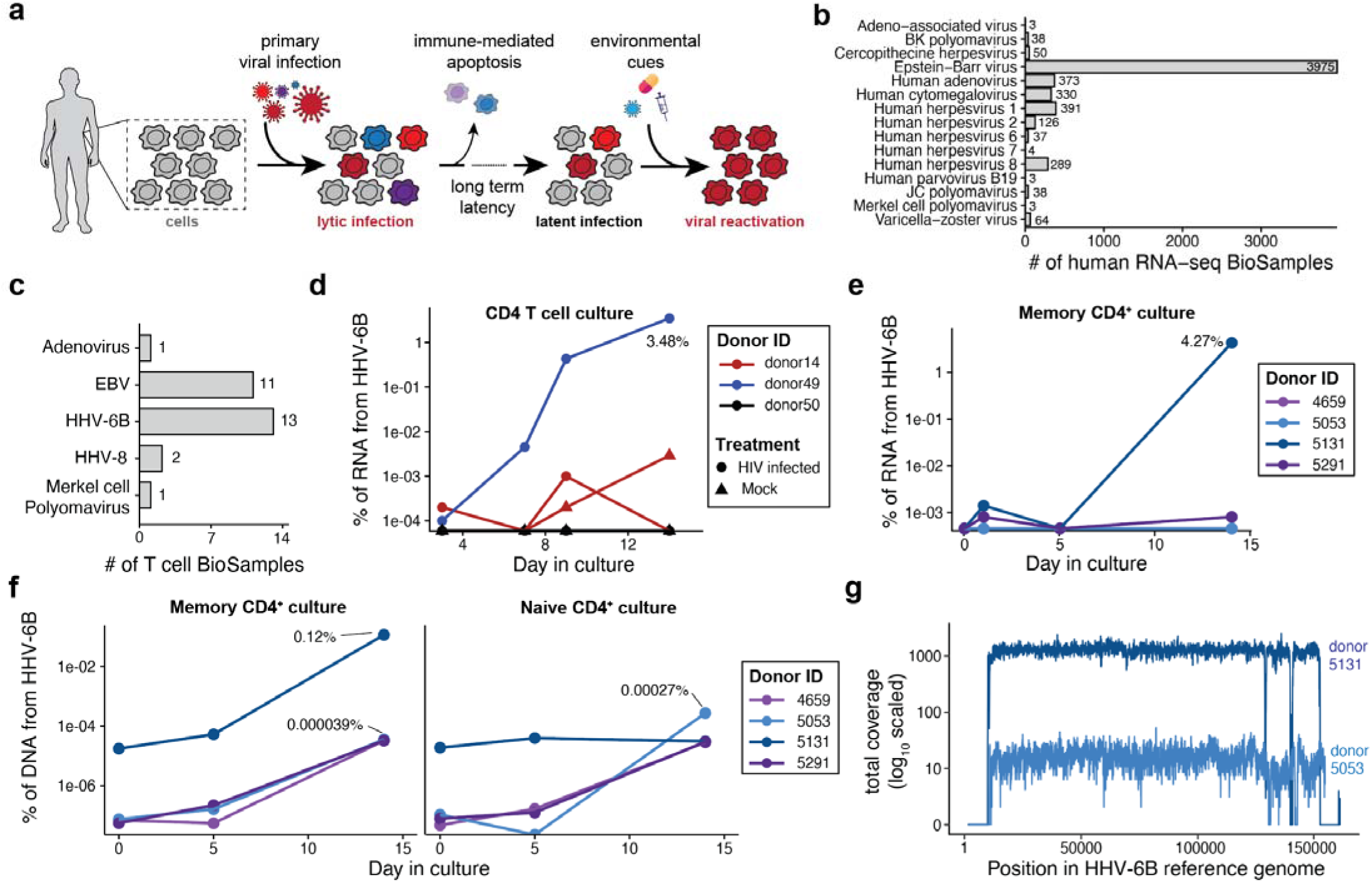
Petabase-scale analysis of viral nucleic acids reveals that HHV-6 is reactivated in human T cells. **(a)** Schematic of viral life cycles of endogenous human viruses. Viruses can enter a latent phase after primary infection and become reactivated based on environmental cues. **(b)** Enumeration of BioSamples with human viruses capable of reactivation confidently expressed in human SRA samples. **(c)** Summary of BioSamples with viral expression in human T cells. Viruses shown in (b) but not (c) had no evidence of reactivation in T cells. **(d)**Reanalysis of Shytaj *et al*.^17^ RNA-seq data. CD4+ T cells from 3 separate donors were treated with either an HIV or mock infection and cultured for ~2 weeks. Shown is the % of RNA molecules aligning to the HHV-6B reference Serratus reference. **(e)** Reanalysis of the LaMere *et al*.^16^ data. Naive and memory CD4+ T cells were separated and cultured for two weeks. Shown is the % of RNA molecules aligning to the HHV-6B reference Serratus reference. **(f)** Quantification of HHV-6B episomal DNA from a reanalysis of ChIP-seq from LaMere *et al*.^16^ data. Shown are the % of DNA reads uniquely mapping to the **H** HV-6B transcriptome, noting the percent of total DNA reads mapping to the HHV-6B reference genome. **(g)** Co verage of the HHV-6B genome across two ChIP-seq libraries from different donors LaMere *et al*.^16^. Repetitive regions on the ends of the chromosome show no uniquely mapping reads.

In an effort to study human viral reactivation systematically, we mined a petabase-scale resource, Serratus^12^, that quantified viral abundance from RNA sequencing data from the Sequence Read Archive (SRA). Here, we considered the occurrence of high-confidence viral RNA from 129 curated human viruses^13^, focusing on the 19 viruses across the *Herpesviridae, Polyomaviridae, Adenoviridae*, and *Parvoviridae* families that are known to have lytic replication cycles and can reactivate^9^ (**Methods**). For 15 of these 19 viruses, we observed a high confidence annotation of viral RNA in at least one BioSample, totaling 5,724 samples with plausible viral RNA expression, and in turn, potential lytic reactivation (**Fig. 1b; Methods**). After defining this landscape, we examined samples with the highest levels of viral RNA, which predominantly encompassed Epstein-Barr Virus (EBV) expression in B cells or intentional over-expression of a specific virus in a model cell line system (**Extended Data Table 1**). Surprisingly, we observed that for human herpesvirus 6 (HHV-6), the top positive sample came from primary T cell culture where no record of the HHV-6 virus was noted in the original study, suggesting that the virus may have been reactivated (**Fig. 1c; Methods**). Further, we observed high levels of HHV-6 expression from various independent studies of T cells with and without *ex vivo* culture (**Extended Data Table 2**). Overall, we observed evidence of 5 total viruses reactivating in human T cell samples (**Fig. 1c;** (**Extended Data Table 2**; **Methods**), with HHV-6B expressing orders of magnitude more viral RNA than other detected viruses (**Fig. 1c;** (**Extended Data Table 2**; **Methods**). Thus, we sought to more closely examine the link between HHV-6 reactivation and primary T cells.

Whereas HHV-6A infections are largely confined to sub-Saharan Africa^14^, the HHV-6B virus accounts for 97-100% of HHV-6 infections reported in the United States, Europe, and Japan. The canonical receptor for HHV-6B, TNFRSF4 (CD134 or OX40), is upregulated during CD4+ T cell activation as well as CD8+ T cell activation to a lesser extent (as well other healthy tissues; **Extended Data Fig. 1, 2**). Thus, the expression of OX40 supports the notion that this viral species could spread through endogenous HHV-6 positive T cells during *in vitro* T cell culture and expansion or *in vivo.^15^* We examined sequencing data from the two studies with high levels of HHV-6^16,17^ that performed longitudinal sampling of cultured T cells in standard conditions (anti-CD3/anti-CD28 bead activation with IL-2) to understand the kinetics of viral reactivation in T cell cultures. First, Shytaj *et al.^17^* cultured total CD4+ T cells for 14 days from three separate donors where cells were initially treated with either HIV or mock infection (we note that transcriptional homology between HIV and HHV-6B could not account for the observed HHV-6B expression; see **Methods**). Here, two donors (14 and 49) showed clear evidence of HHV-6B reactivation over the 2 weeks in culture, including a sample where nearly 3.5% of all RNA molecules were derived from HHV-6B and mapped to virtually all of the HHV-6B transcripts (**Fig. 1d; Extended Data Fig. 3a**). Separately, analysis of the LaMere *et al.^16^* study, in which naive and memory CD4+ T cells were separated and cultured for two weeks before profiling with RNA-seq (**Fig. 1e; Extended Data Fig. 3b**) and chromatin immunoprecipitation with sequencing (ChIP-seq; **Fig. 1f,g**) demonstrated that one donor expressed HHV-6B transcripts at high levels (~4.27% of all RNAs) after 2 weeks of memory CD4+ T cell culture *in vitro*. As this study utilized the same cell cultures to perform ChIP-seq against a variety of histone modifications, we remapped sequencing reads to estimate the abundance of HHV-6 derived DNA from these samples (**Methods**). We observed that all three donors increased their HHV-6B DNA copy number over the duration of culture, including a sample from donor 5131 that had the highest viral copy number for both DNA and RNA (**Fig. 1e,f; Methods**). We confirmed that these mapped DNA molecules were derived from the HHV-6B virus, noting the uniform coverage across the HHV-6B reference genome outside repeat regions (**Fig. 1g; Methods**) and across all transcripts. Finally, while our petabase-scale mining identified HHV-6 as a latent virus that may be reactivated in T cells *in vitro*, we further observed BioSamples with high HHV-6B derived from patients with bronchiolitis^18^, graft versus host disease (GvHD)^19^, and purified T cell populations from patients with cutaneous T-cell lymphoma (CTCL)^20^ (**Extended Data Fig. 3c,d**). In summary, analysis of these public datasets implicates the reactivation and lytic activity of HHV-6B in T cells *in vitro* and *in vivo*.

In light of eight cases of HHV-6 encephalitis from patients receiving CAR-T cell therapy, including in three clinical trials^1,5–7^, we hypothesized that the lytic HHV-6 may be derived from cell therapy products. Using research grade allogeneic CAR-T cell culture conditions, we cultured cells over a 19-day period and screened cells with qPCR to detect the U31 gene from the HHV-6B virus (**Methods**). Over the course of screening, we identified four exemplar PBMC lots derived from healthy donors (**Methods**) that by day 19 expressed a range of HHV-6B viral DNA from 0.0015-0.90 copies of U31 per cell (**Fig. 2a**). For two of these donors, we also conducted longitudinal qPCR assays on the cultures, which demonstrated low, if any, early HHV-6 DNA before a rapid increase after approximately two weeks in culture (**Fig. 2b**). These data directly support our observations from the Serratus analysis that HHV-6 nucleic acid can become more abundant in T cell cultures over time.

**Figure 2.**
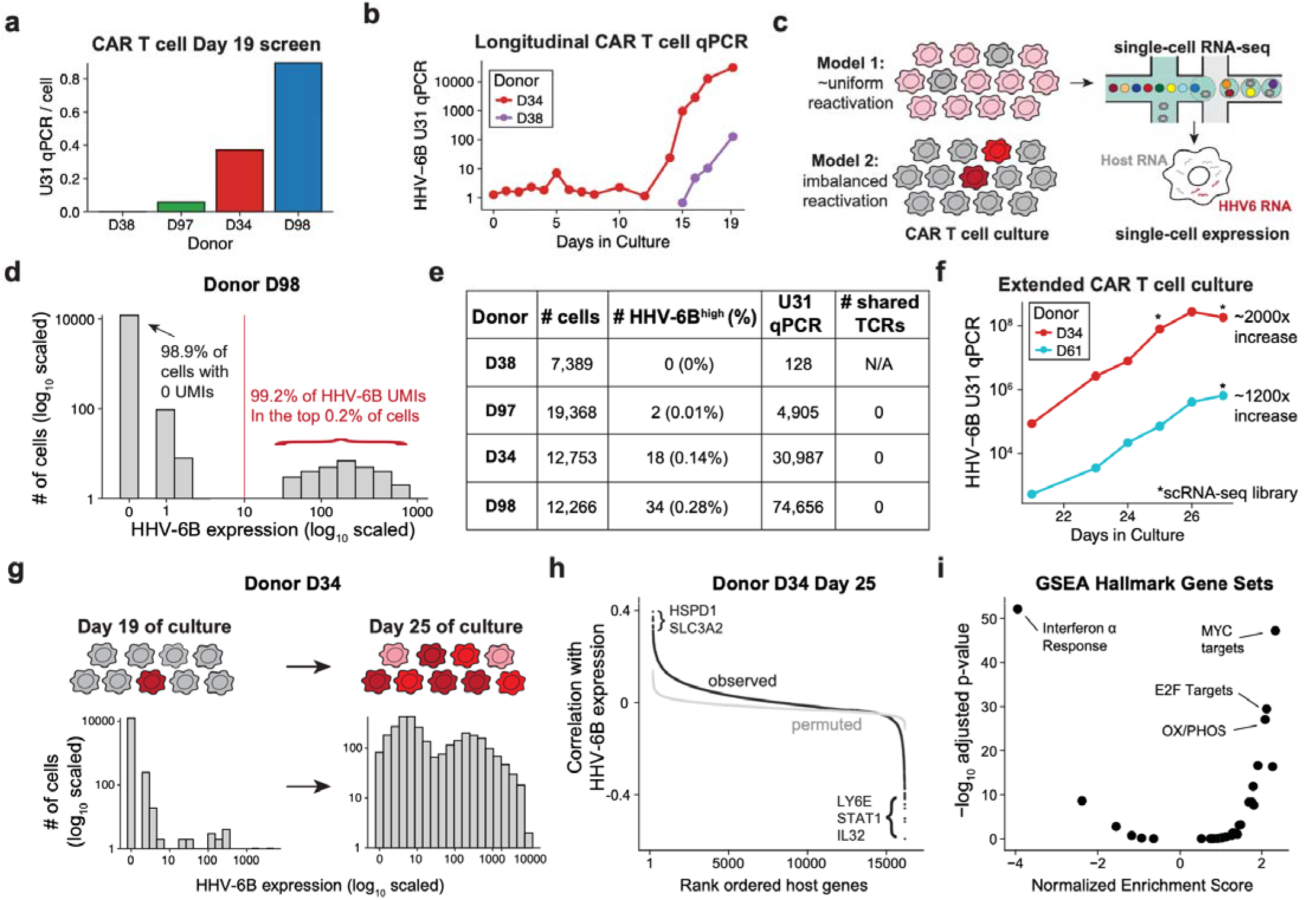
scRNA-seq identifies rare HHV-6 expressing cells in CAR-T cell culture. **a)** Summary of HHV-6B qPCR expression at Day 19 for four donors normalized to the total cell count. Bars are sorted in increasing order of expression. **(b)** Longitudinal sampling of the HHV-6B U31 gene copies CAR-T cell culture from two donors. **(c)** Schematic of single-cell sequencing workflow to detect HHV-6^+^ cells from the CAR-T culture. Two models are presented that would explain HHV-6 reactivation: Model 1 (top) where cells all express HHV-6 transcripts or Model 2 (bottom) where only a subset of cells express HHV-6B. Both the host and HHV-6B viral RNA can be directly quantified using our 10x Genomics RNA-seq workflow. **(d)** Summary of HHV-6B expression from an individual donor (D98). The top 0.2% of cells contain 99% of the HHV-6B transcript UMIs from this experiment. **(e)** Tabulated summary of scRNA-seq profiling for four CAR-T donors, including number of cells profiled, % expressing HHV-6, U31 qPCR value, and number of shared TCR clones between the HHV-6B^+^ cells. **(f)** Extended longitudinal sampling of the HHV-6B U31 qPCR CAR-T cell culture for two donors. The fold increases from the first qPCR measurement (Day 21) and the last (Day 27) are noted for each donor. **(g)** Schematic and summary of HHV-6B expression in the D34 donor after 19 and 25 days, showing evidence of HHV-6B spreading in the culture as depicted in the schematic. **(h)** Correlation analyses of host factor gene expression with HHV-6B expression in individual cells. The per-gene correlation statistics are shown in black against a permutation of the HHV-6B expression in gray. Noteworthy genes are indicated. **(i)** Pathway enrichment analysis of GSEA/Msigdb Hallmark gene sets. A positive Normalized Enrichment Score (NES) corresponds to genes that are overexpressed in cells with high amounts of HHV-6B transcript.

**Figure 3.**
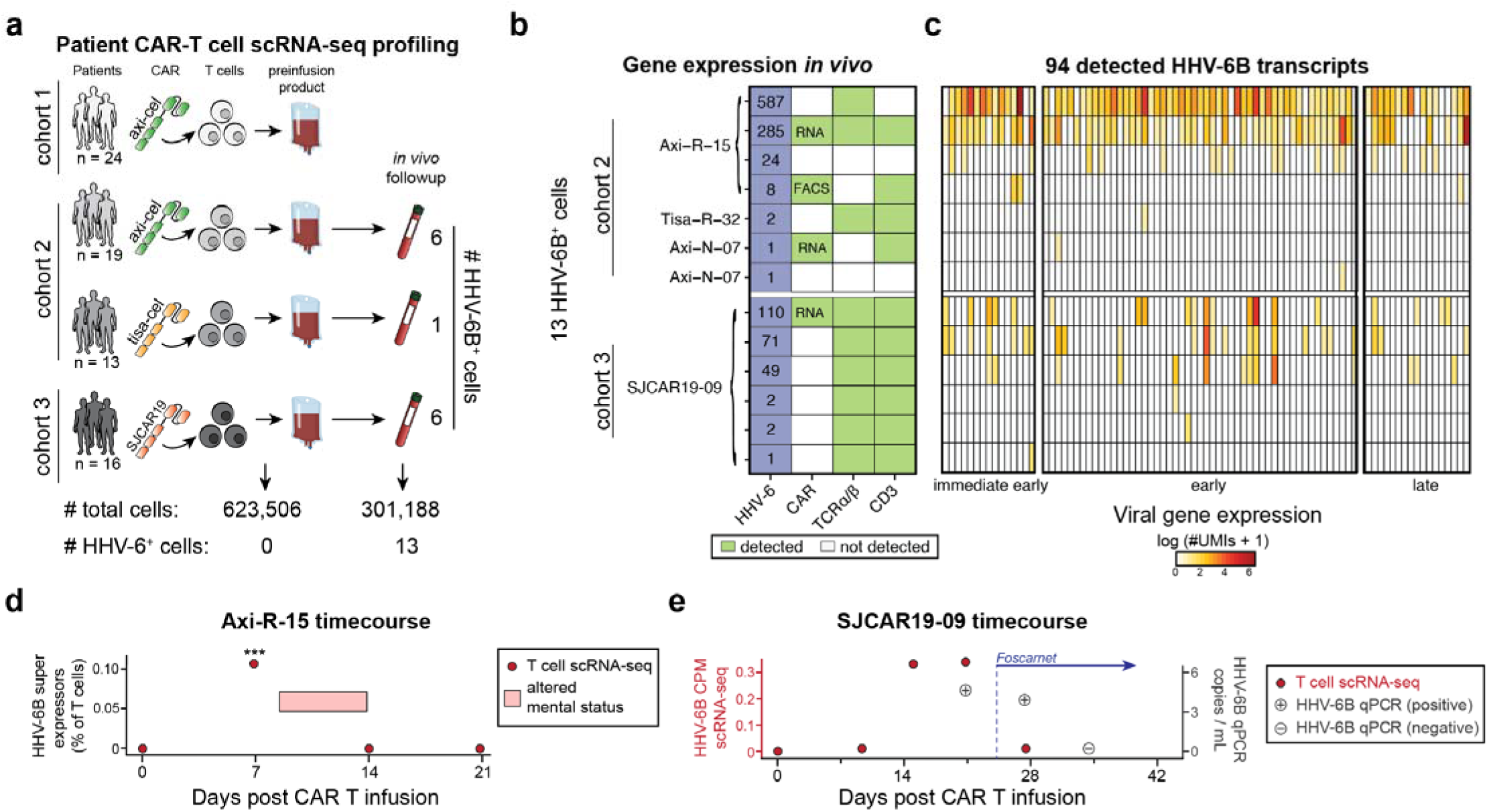
FDA-approved CAR-T cells and CAR-T cells from clinical studies reactivate HHV-6 *in vivo*. **(a)** Schematic and summary of patient cohorts, noting different cell therapy products (axi-cel and tisa-cel) and two different timepoints for sampling (pre-infusion product and weeks post-infusion). The number of total cells analyzed and HHV-6+ cells detected are noted underneath. **(b)** Summary heatmap of 7 HHV-6 positive cells from cohort 2 *in vivo* follow up and 6 HHV-6 positive cells from cohort 3 post infusion samples. Number of HHV-6 transcripts is noted in blue boxes; non-zero host expressions are noted in green boxes (see **Methods**). **(c)** Heatmap of HHV-6B transcripts (columns) by same individual cells (rows, as in **(b)**) grouped by viral gene class (immediate early; early; late expressing genes as previously described^27^). **(d)** Summary of Axi-R-15 patient, noting 4 libraries of scRNA-seq (red dot) and the clinical period corresponding to altered mental status. Stars represent statistical significance of two-sided binomial tests (day 7 vs. day 0 *p* = 0.00047; day 7 vs. day 14 *p* = 0.00032). **(e)** Summary of SJCAR19-09 patients, including PCR of HHV-6 and T cell scRNA-seq abundance. Treatment regimen with Foscarnet is noted with initial dose administered at day +24.

To better understand the dynamics and cell states of reactivated HHV-6B during these CAR-T cell cultures, we considered whether HHV-6 would be uniformly reactivated in positive samples (Model 1) or enriched in only a subset of cells that spread the virus throughout the culture (Model 2; **Fig. 2c**). Consistent with recent efforts that quantify both viral and host gene expression at a single-cell resolution^21^, we established an efficient pseudoalignment pipeline after single-cell RNA-seq (scRNA-seq) to quantify HHV-6B transcripts with no detectable false positive quantification (**Methods**). Strikingly, our single-cell characterization of HHV-6+ CAR-T cell cultures revealed the presence of rare HHV-6+ cells that we term ‘super-expressors’, directly supporting Model 2 where only a rare subset of cells initially express HHV-6B (**Fig. 2c; Extended Data Fig. 4a**). We detected expression of 90 HHV-6B transcripts, including immediate early (IE) genes that were enriched among a subset of super-expressors, suggesting that these cells had recently been infected or had reactivated HHV-6B (**Extended Data Fig. 4b**). In total, these rare super-expressors accounted for only 0.2% of total cells in culture for donor D98 despite expressing a vast majority of the total HHV-6 viral RNA detected (99.2%; **Fig. 2d**). Across three donors (D34, D97, D98), we confirmed the existence of this rare cell population as we identified between 0.01%-0.3% (or 1 in 360-10,000) of cells that reactivate or express HHV-6 transcripts at high levels. Utilizing the host (T cell) gene expression signatures, we performed standard dimensionality reduction and annotation, revealing that super-expressors were confined to the CD4+ T cell compartment, consistent with relative over-expression of the HHV-6B receptor OX40 expression in T cell subsets (**Extended Data Fig. 1a; Extended Data Fig. 4c; Methods**). We note a fourth donor (D38) had no detectable HHV-6B^hi^ cells, corresponding to the lowest U31 qPCR value (donor D38) (**Fig. 2e**). Sequencing of TCR α/ß chain transcripts revealed that none of the HHV-6B+ cells from these donors shared a clonotype receptor sequence (**Fig. 2e**). As we could quantify both viral and host gene expression in single-cells, we could further link host genes with HHV-6B expression (**Methods**). Differential gene expression of HHV-6 super-expressors compared to other HHV-6B-cells showed a consistent up-regulation of lymphotoxin α (LTα) and downregulation of lymphotoxin β (LTβ; **Extended Data Fig. 4d; Methods**), but we were unable to nominate host factors, such as surface markers, that would clearly segregate cells capable of HHV-6 reactivation from other cells.

For two donors, D34 and D61, we extended the time of T cell expansion and collected scRNA-seq libraries after a total of 25 and/or 27 total days from the initiation of the culture. Analysis of these later time points revealed an increase in the abundance of HHV-6B transcripts by ~1,200-2,000-fold, compared to day 20 (**Fig. 2f**). At these later timepoints, we determined that 49% and 62% of all T cells were HHV-6 super-expressors, indicating that the lytic reactivation of the virus spread rapidly and suggesting that HHV-6 can rapidly spread and overtake certain T cell cultures (**Fig. 2g; Extended Data Fig. 4e**). In these samples with greater numbers of HHV-6B+ cells, we utilized a correlation statistic to identify hundreds of genes associated with HHV-6B infection following reactivation in culture (**Fig. 2h**, **Extended Data Table 3**). Of note, this analysis identified proviral genes like *HSPD1* and *SLC3A2*, that have been characterized as positive regulators of other infections^22,23^ (**Fig. 2h**). Conversely, we identified an upregulation of genes such as *LY6E^24^, STAT1^25^*, and *IL-32^26^* in cells low for HHV-6B, each of which has previously been implicated in antiviral control for other diverse infection contexts (**Fig. 2h**). Further, after stratifying the HHV-6B transcripts into IE, early, and late using a prior annotation^27^, the only gene signature correlated with OX40 expression in single-cells was the IE signature, aligning with the expectation that recently infected cells would express higher levels of the viral receptor (**Extended Data Fig. 4f; Methods**). Integrated analyses of altered gene expression signatures associated with HHV-6B expression levels corroborated the role of various pro- and antiviral pathways, including type I interferon response^28^ and oxidative phosphorylation^29^ (**Fig. 2i; Methods**). Collectively, our host-viral association analyses indicate that a subset of CAR-T cells in culture that resist infection are transcriptionally inducing an anti-viral response, spanning metabolic transporters, transcription factors, cytokines, and surface molecules.

Having characterized this rare HHV-6+ population in T cell culture, we hypothesized that these cells could be a potential source of HHV-6 encephalitis previously reported in CAR-T trials and case studies^1,5–7^. To test this, we reanalyzed three cohorts of scRNA-seq datasets collected from B cell lymphoma and leukemia patients receiving one of three CAR-T cell products, including U.S. Food and Drug Administration (FDA) approved axicabtagene ciloleucel (axi-cel) and tisagenlecleucel (tisa-cel), as well as from one early phase clinical study (SJCAR19; NCT03573700). We note that all CAR-T cells were engineered to target the CD19 antigen. Across these three cohorts, we screened single-cell expression profiles of 924,694 cells from 72 patients for HHV-6B expression, including T cells from the pre-infusion products and cells from peripheral blood draws weeks after the infusion (**Fig. 3a; Table 1**). While we did not detect any HHV-6+ cells in the preinfusion products, we detected a total of 13 cells from 4 samples that expressed 1 or more HHV-6B transcripts in the follow-up blood draws, including eight ‘super-expressor’-like cells (**Fig. 3b;** see **Methods** for discussion of cells with 1 or 2 UMIs). Through a combination of CAR protein, CAR transcript, TCR α/β transcripts, and CD3 expression, we verified the cellular identity of 29 total HHV-6B+ cells as engineered T cells (**Fig. 3b; Extended Data Table 4; Methods**). We additionally confirmed that these cells expressed a diversity of HHV-6B-associated transcripts among these *in vivo* super expressors (**Fig. 3c**). Thus, having confirmed the presence of HHV-6B+, CAR+ T cells *in vivo*, we sought to examine the course of the two patients with detectable super-expressors from the post-infusion blood draws.

**Table 1.** Summary of HHV-6B expression in large-scale single-cell datasets. SE = HHV-6 super expressors.

| Description | # Cells | # Reads | #HHV-6B UMIs | #HHV-6+ cells (# SE) | Ref. |
| --- | --- | --- | --- | --- | --- |
| 24 patient CAR-T cell pre-infusion product | 137,326 | 8,972,716,245 | 0 | 0 (0) | 36 |
| 32 patient (CAR) T cell, pre- and post- infusion | 602,577 | 15,403,016,654 | 919 | 7 (4) | 2 |
| 16 patient (CAR) T cell, pre- and post- infusion | 184,791 | 81,875,146,341 | 374 | 6 (3) | 3 |
| 1 patient (axi-R-15) pre- and post- infusion followup | 31,953 | 3,086,842,082 | 789 | 15 (6) | This Study |
| Human cell landscape, 60 tissue types | 599,926 | 15,680,467,351 | 0 | 0 (0) | 31 |
| ~1,000 donor PBMCs | 1,449,385 | 53,872,337,003 | 6 | 0 (0) | 37 |

First, we examined samples from the subject, axi-R-15, with cells showing the highest number of captured HHV-6B transcripts. We complemented the existing scRNA-seq profiles by sequencing another 31,953 cells for this subject spanning four timepoints, including weekly blood draws out to day +21 (**Fig. 3d; Methods**). Along this time course, we observed HHV-6+ cells only at day 7 where the HHV6+ super-expressors were present with an overall frequency of ~1 in 1,000 T cells (a total of 10 super-expressors were detected among 9,757 T cell scRNA-seq profiles). Notably, this patient was clinically diagnosed with immune effector associated neurotoxicity syndrome (ICANS) following axi-cel treatment and subsequently developed altered mental status with transient hypotension beginning on day +9, which peaked on day +12 (grade 3a by CTCAE criteria, grade 2 by ASTCT) before returning to baseline by day +14 (**Fig. 3d**). Similarly, we observed a spike in HHV-6+ cells at day +7 that was resolved by day +14. While we lack direct evidence that the reactivated HHV-6 in CAR-T cells caused the altered mental state observed in this patient, we note the dynamics of viral reactivation and clearance could be observed within 1 week of each other (**Fig. 3d**).

Second, we detected three HHV-6 super-expressors in cohort 3 from patient SJCAR19-09. Though two patients were tested for HHV-6 via qPCR from this cohort, only SJCAR19-09 tested positive (**Fig. 3e**), corroborating our detection of HHV6+ CAR-T cells in this individual. SJCAR19-09 tested positive via qPCR at day +21 post-CAR-T cell infusion and was started on Foscarnet at day +24 (**Fig. 3e**). After another positive qPCR test on day +27, the patient no longer had detectable levels of HHV-6 at day +34 (**Fig. 3e**). Similarly, we detected HHV6+ super-expressors at day +14 that persisted *in vivo* until day +21 before these cells were undetectable at day 27. As the products used in cohorts 2 and 3 were cultured for 7-10 days before infusion, we speculate that HHV-6 may become reactivated in some cells after the therapy is administered but the virus was not yet transcriptionally active or sufficiently abundant to be detected via single-cell sequencing. However, after 1-2 weeks of persistence and expansion *in vivo*, these rare HHV6+ super expressors were detectable in peripheral blood draws. Importantly, the SJCAR19-09 case vignette suggests that Foscarnet may be an efficacious agent to mitigate the effects of viral infection during the course of therapy if HHV-6 expressing cells are identified during the course of treatment with CAR-T cells.

Though the population of HHV-6+ CAR-T cells from these patients is rare, these results directly link CAR-T cell therapy to a potential cellular reservoir of lytic HHV-6 *in vivo*. Nevertheless, we sought to screen other possible cellular sources of viral expression throughout human tissues and estimated HHV-6B transcript abundance from large consortiums profiled with scRNA-seq (**Table 1**). First, we confirmed that among ~1,000 peripheral blood mononuclear cell (PBMC) samples, each stemming from a different donor, we did not detect any HHV-6+ cells (**Table 1**) or from the apheresis sequencing samples from our previous CAR-T datasets. These analyses suggest that HHV-6 is not transcriptionally active in peripheral blood under normal circumstances and unlikely to be the source of HHV-6 without the T cell therapy culture *in vitro*, albeit for a period of time that generally exceeds the manufacturing time in CAR-T cell therapy. We draw the distinction that while a majority of individuals in North America (where most of the T cells were sourced for these studies) are seropositive for prior HHV-6 infection and often have detectable viral DNA from PCR^30^, our results indicate that transcriptionally active virus is a rare phenomenon in peripheral blood of adults. Therefore, measures that ensure lack of transcriptional HHV6 activity in the CAR-T infusion product are warranted to prevent infusion related HHV6 infection. Second, we quantified the human cell landscape^31^, consisting of >700,000 cells from 60 tissue types (**Methods**). We did not detect a single HHV-6B transcript in this dataset (**Table 1**). Though a variety of epithelial cells express the OX40 receptor (**Extended Data Fig. 2**), this negative result further supports the possibility that lytic HHV-6 in patients with encephalitis originates from the cellular therapy product.

As HHV-6 reactivation has been recurrently observed in other settings, notably bone marrow and HSCT^32^, we cannot exclude other cellular sources of pathogenic HHV-6 as both cell therapies are introduced into an immunosuppressed host. Additionally, in settings where lytic HHV-6 is introduced into the body from the CAR-T cells, the physiological implications of this infection are not straightforward. From our pan-tissue gene expression analyses, we observed the expression of OX40 on endothelial cells lining the blood-brain barrier (**Extended Data Fig. 2b**), and that OX40 expression is induced in endothelial cell lines in the presence of TNF-α, a cytokine typically circulating during CAR-T therapy (**Extended Data Fig. 2c**). These data suggest the possibility that HHV-6 could more easily infect these blood-brain endothelial cells during latent reactivation, thereby contributing to viral encephalitis. Although further experiments are required to establish this possibility, we speculate that the lytic virus from HHV-6+ reactivation may infect these brain endothelial cells to contribute to viral-induced encephalitis and confusion as previously reported in a CAR-T treated patient^1^. We note however that the diagnosis of HHV-6 encephalitis in the setting of CAR-T is part of the differential diagnosis of CAR-T related ICANS, and that a firm diagnosis of HHV-6 encephalitis is not always distinguishable from ICANS, even in the presence of HHV-6 viremia.

It is possible that latent viral reactivation may extend beyond HHV-6 and T cells for other viruses or forms of cell therapy. From our inferences mining the Serratus database, we observed reactivation of HSV-1 from reprogrammed induced pluripotent stem cells (iPSCs) in 18 RNA-seq libraries (**Extended Data Table 5**) as well as EBV expression in T cells (**Fig. 1c**; **Extended Data Table 2**). We note that the EBV receptor, CD21, is expressed in gamma-delta T cells at levels exceeding B cell subsets in resting and stimulated conditions (**Extended Data Fig. 1b**), leading us to note the possibility that EBV may become reactivated in gamma-delta T cell therapies. Of note, our screen did not nominate a reactivation candidate virus in natural killer (NK) cells. Regardless, we suggest that NK cell therapies may warrant further investigation for potential viral latency as longitudinal cultures were more commonly profiled for T cells and iPSCs than NK cells in the Serratus database, potentially biasing our inferences for this population.

Our work has implications for current and future applications of cell therapies. While we and others have characterized modes of antigen-dependent toxicity^33–35^, we implicate the T cells as a potential source of a lytic virus, HHV-6, in a presumed antigen-independent mode of toxicity. We observed HHV-6 reactivation in both native (**Fig. 1**) and engineered T cell states (**Fig. 2**), including FDA-approved CAR-T cell products and CAR-T cells from another clinical study (**Fig. 3**). Altogether, this suggests that HHV-6 reactivation occurs in a CAR-independent capacity and instead may be linked to T cell activation and proliferation. We observed the increase of lytic HHV-6 over time both *in vitro* (**Fig. 2**) and *in vivo* (**Fig. 3**), indicating that screening the cell therapy product for live virus at a single time point (e.g., pre-infusion product) may not be fully adequate to eliminate virus-positive cells from the final therapeutic product, although longer culture times may improve the sensitivity of screening prior to infusion. Indeed, we determined that while HHV-6 may initially be transcriptionally active in a subset of cells, these rare cells can infect the remaining cells in culture (**Fig. 2**). In an immunocompromised host, we note the possibility of similar spreading occurring *in vivo*. However, in the two cases where we observed HHV-6+ cells *in vivo* (**Fig. 3**), we observed clearance of the HHV-6 infected cells within 1-3 weeks for both donors. In sum, we suggest that longitudinal viral screening may be warranted in patients receiving cell therapies, and that a suitable control strategy for CAR-T drug product should include transcriptome analysis to demonstrate absence of HHV-6 replication. While we advocate for closely monitoring patients for any potential toxicities, this work supports routine assessment of HHV-6 expression in patients, particularly in the setting of clinical symptoms such as unexplained fevers or neurotoxicity. In total, our efforts demonstrate the utility of comprehensive genomics sequencing and analyses to disentangle complex clinical phenotypes and nominate an unappreciated route of viral infection via cell therapy.

## Methods

### Serratus viral screen analyses

To examine human viruses that may become transcriptionally activated in a systematic approach, we queried the Serratus^12^ database for 129 human viruses curated from ViralZone^13^ using the NC genome identification provided on the ViralZone webpage as input in the Serratus API. To focus on viral reactivation, we subset to 19 viruses across the *Herpesviridae, Polyomaviridae, Adenoviridae*, and *Parvoviridae* families contained in the query. We categorized a sample as a potential hit (**Fig. 1b**) given the Serratus output returned a minimum of 10 mapping reads with a homology score of at least 90% for the specific BioSample/virus pair.

As the ViralZone genome annotation for HHV-6 corresponded to HHV-6A (NC_001664) is the dominant species in sub-Saharan Africa^14^, we queried the Serratus database for HHV-6B (NC_000898) for the T cell specific analyses shown in **Fig. 1c-e**. To annotate BioSamples as belonging to specific classes, we performed a regular expression mapping for both the GEO sample title and the description. For T cells, our regular expression was “T-cell|\ T\ cell|Tcell|CD4T|CD8T|^^^T cell” whereas for iPSCs, we queried “iPSCļluripotent”. Notably, we did observe expression of EBV from annotated T cell populations (**Fig. 1c**), including intestinal intraepithelial cytotoxic CD8 T cells.^38^ As EBV infects cells via CD21, a canonical B cell marker, we examined the expression of CD21 across all resting and stimulated immune cell subsets, revealing that gamma-delta T cells can express CD21 at higher levels than any B cell subset (**Extended Data Fig. 1b**), leading us to speculate that EBV reactivation may be a possibility in gamma-delta T cells. Overall, we also suggest that the systematic approach pairing viral expression to specific BioSamples via Serratus could enable the identification of cell types to become infected that have not been previously characterized.

We note that the two primary studies that showed HHV-6 reactivation in T cell culture, Shytaj *et al.^17^* and LaMere *et al.^16^*, took place on different continents, making it highly unlikely that other factors enabled the reactivation of HHV-6 in their cultures (e.g. same donor) aside from T cell culture induced HHV-6 reactivation. For our deeper analyses of these datasets (**Fig. 1d,e**), we paired the Serratus read abundance with metadata from GEO, including the total number of reads per library. In the Shytag et al. dataset, cells were treated with an HIV infection. We reanalyzed the gene expression from this data at the per-gene level, verifying a diversity of transcripts were expressed and regions of high homology between HHV-6B and HIV were not the only transcripts detected (also noting that not all HIV-infected cells showed HHV-6B expression). Thus, the inference of HHV-6B reactivation in this setting is not attributable to off-target viral read mapping.

For ATAC-seq and ChIP-seq libraries, DNA was not processed via the Serratus workflow, requiring realignment to the HHV-6B reference genome (e.g. **Fig. 1 f,g**). Thus, we downloaded raw .fastq files from GEO and aligned reads to the HHV-6B reference genome (Genbank AF157706) using bwa and retaining only uniquely mapped reads (MAPQ >30) for downstream analysis and quantification. To screen for reads that were outside the repetitive regions (which have high homology with regions of the human genome), we required the read to map within the (8788, 153321) coordinates.

### Primary human T cell cultures

Peripheral blood mononuclear cells (PBMCs) were isolated from leukopaks obtained from healthy donors and stored in liquid nitrogen prior to use. PBMCs were thawed, resuspended in X-vivo media (Lonza) supplemented with 5% human serum (Access Cell Culture) and IL-2 (Miltenyi), and stimulated with T Cell TransAct (Miltenyi). Cells were transduced with a lentivirus containing proprietary CAR sequences early in culture followed by further genetic manipulation for immune evasion and prevention of graft rejection. These T cell cultures were allowed to expand for up to 19 days and then cryopreserved using a controlled rate freezer prior to storage in liquid nitrogen. For selected donor PBMCs (**Fig. 2f**), the 19 day cryovial was thawed and recultured and expanded up to 7 additional days. PBMC donor lots were screened at Allogene Therapeutics for HHV-6 expression following these CAR-T cell culture conditions noted herein. A subset of the lots screened included in the manuscript (**Fig. 2**) and were included to demonstrate a range of HHV-6B expression.

### U31 qPCR protocol

Primary human T cells were centrifuged at 540x *g* for 5 min before removal of culture supernatant, leaving the cell pellet for quantitative polymerase chain reaction (qPCR). Cells were processed for DNA using the manufacturer’s protocol from Qiagen DNeasy Blood & Tissue Kit (Qiagen). Concentration and 260/280 ratio of the final product was measured using a NanoDrop Spectrophotometer (ThermoFisher).

For qPCR, the master mix used was TaqMan Fast Virus 1-Step Master Mix (ThermoFisher). Manufacturer’s protocol was followed to add the appropriate volumes of master mix, water, DNA and primers. All samples were prepared in triplicate and run on a QuantStudio 6/7 Flex Real-Time PCR System (ThermoFisher). Primer sequences for designated HHV-6 marker (U31) were designed based on prior reports^39^ and were modified as follows: FW seq: 5’-CGACTCTCACCCTACTGAACGA-3’; RV seq: 5’-GAGGCTGGCGTCGTAGTAGAA-3’; Probe seq: 5’-/5YakYel/AGCCACAGC/ZEN/AGCCATCTACATCTGTCAA/3IABkFQ/-3’. Measurements were taken in triplicates and the mean value is shown for qPCR plots.

### Single-cell RNA-seq

Libraries for scRNA-seq were generated using the 10x Chromium Controller and the Chromium Single Cell 5’ Library Construction Kit and human B cell and T cell V(D)J enrichment kit according to the manufacturer’s instructions. Briefly, the suspended cells were loaded on a Chromium controller Single-Cell Instrument to generate single-cell Gel Bead-In-Emulsions (GEMs) followed by reverse transcription and sample indexing using a C1000 Touch Thermal cycler with 96-Deep Well Reaction Module (BioRad). After breaking the GEMs, the barcoded cDNA was purified and amplified, followed by fragmenting, A-tailing and ligation with adaptors. Finally, PCR amplification was performed to enable sample indexing and enrichment of scRNA-Seq libraries. For T cell receptor sequencing, target enrichment from cDNA was conducted according to the manufacturer’s instructions. The final libraries were quantified using a Qubit dsDNA HS Assay kit (Invitrogen) and a High Sensitivity DNA chip run on a Bioanalyzer 2100 system (Agilent). 10x scRNA-seq libraries were sequenced as recommended by the manufacturer (~20,000 reads / cell) via a Nova-seq 6000 using an S4 flow cell. Raw sequencing data was demultiplexed using CellRanger mkfastq and aligned to the host reference genome using CellRanger v6.0 and TCR sequences were processed using the CellRanger vdj pipeline with default settings.

### Single-cell HHV-6B quantification workflow

To quantify HHV-6B reads, we sought to develop a workflow that a) would utilize pseudoalignment rather than standard transcriptomic alignment, b) could be mapped to only the HHV-6B reference transcriptome (rather than a mixed transcriptome), and c) had no detectable off-target expression (i.e. 0 reported HHV-6B expression for scRNA-seq samples from human transcripts). Our rationale for these constraints was to optimize computational efficiency in order to reprocess massive-scale scRNA-seq libraries, noting ~94 billion paired end reads were processed in the studies noted in **Table 1**.

To satisfy developed a kallistoļbustools^40,41^ workflow to rapidly quantify reads from either single-cell or bulk transcriptomes. We downloaded the GenBank AF157706 reference transcriptome and created a kallisto index using the default -k 31 (kmer) parameter. For single-cell libraries, raw sequencing reads were processed using the kallisto ‘bus’ command with appropriate hyperparameters for each version of the single-cell chemistry (either 14 or 16 bp sequence barcode and 10 or 12 bases of UMI sequence). After barcode and UMI correction, a plain text sparse matrix was emitted, corresponding to unique HHV-6B reads mapping to individual cells in the single-cell sequencing library. For bulk libraries, the same index could be utilized with the standard kallisto ‘quant’ execution.

To assess our specificity (i.e. no human transcript expression being reported as an HHV-6B transcript), we quantified 4 healthy PBMC libraries that had no expected HHV-6B expression. We recurrently observed non-zero expression at only the AF157706.1_cds_AAD49614.1_1 and AF157706.1_cds_AAD49682.1_98 coding features, both genic transcripts that encode for the DR1 gene, which corresponded to AF157706 equivalence classes 6 and 120 in the kallisto index. By examining reads that mapped to both HHV-6B transcriptome and the human genome, we determined that a region of the DR1 gene is highly homologous to the human gene KDM2A, which is broadly expressed across tissues, including lymphocytes, leading to ~1-3 reads per million errantly aligning to the HHV-6B transcriptome. By excluding these two equivalence classes, we consistently observed exactly 0 HHV-6B UMIs across libraries derived from peripheral blood mononuclear cells^37^ and across 60 healthy tissue types from various donors.^31^ The number of UMIs per cell shown in **Fig. 2** and **Fig. 3** excludes these potential host-derived genes annotated as viral expression. Finally, verification of the *in vitro* expression of the CAR transgene was confirmed via single-cell quantification of the woodchuck hepatitis virus post-transcriptional regulatory element (WPRE) in all sequencing libraries and in single-cells.

To assign whether a cell is a super expressor, we used a semi-arbitrary cutoff of 10 HHV-6B UMIs per cell based on empirical expression (e.g. **Fig. 2d**) and the ease of use/interpretation of the number 10. We note that one cell in the *in vivo* dataset expressed 8 UMIs (**Fig. 3b**), which likely is a super expressor but under sequenced based on the number of overall UMIs detected for this cell (**Extended Data Table 4**). For visualization across the Day 19 samples (**Extended Data Fig. 4a**), we show both the rank-ordered expression per cell based on total HHV-6B UMIs as well as a null model where HHV-6B reads are allocated proportional to the number of total UMIs detected for that particular cell barcode. For HHV-6B transcript heatmaps (**Extended Data Fig. 4b; Fig. 3c)**, we utilize the per-gene annotation from a prior report.^27^ For simplicity in viewing expression signatures, we collapsed the “intermediate early early” and “immediate early early” into a single gene set for visualization in these two heatmaps.

### Integrated viral / host gene expression analyses

To determine differentially expressed genes between HHV-6+ cells and HHV-6^-^ cells from the Day 19 time points, we considered D34 and D38, which had the highest number of HHV-6+ cells. We achieved this using the ‘FindMarkers’ function from Seurat after segregating HHV-6+ cells as those that have a minimum of 2 UMIs (to increase power) and HHV-6^-^ as those with identically 0 HHV-6 UMIs. From the FindMarkers output, we computed a statistic, -log10(p.val)*log2FC, that preserves the direction of over/under expression while partially scaling the gene expression difference by the magnitude of the statistical association, resulting in the plot (**Extended Data Fig. 4d**). From this association, the lymphotoxin genes LTA and LTB were distinguished as the consistent associations As the homotrimer LTα_1_β_2_ is secreted but the heterotrimer (LTcnfc) is membranebound^42^, this suggests that changes in the LT expression upon HHV-6 reactivation lead to a secreted signal via LTα_3_, but due to the pleiotropic effects of this cytokine trimer^42^, the impact of its possible increased secretion is not evident.

To more comprehensively examine host gene association with HHV-6B transcript abundance, we also computed Pearson correlation statistics per-gene performed pathway analyses using the Pearson correlation between the log transformed total HHV-6B UMIs with the Seurat-normalized host expression as a means of rank-ordering genes linked to HHV-6B activity. This was performed for all three re-cultured samples. After broadly verifying the top genes, we presented the data for D34 at day 25 total of culture as it was the earliest time point. Using this correlation-based approach, we identified a significant enrichment of genes downstream of type I interferon activity as being more expressed in cells low for HHV-6B, consistent with the antiviral properties of this pathway^28^. In contrast, targets of the E2F transcription factor were upregulated as the HHV-6B infection spread throughout culture, consistent with prior reports of E2F1-induced HHV-6A expression in T cells^43^. Furthermore, we observed an increase in genes associated with oxidative phosphorylation, a pathway previously reported to be co-opted during active viral infection^29^. Overall, our results largely recapitulate known pro and antiviral responses, indicating that the host transcriptome at the single-cell level permits or resists HHV-6B infection from spreading within the T cell culture.

### Reanalysis of public single-cell data

The data presented in **Table 1** represents a summation of four large datasets reanalyzed for HHV-6B expression. In brief, the number of cells and raw sequencing reads analyzed are reported, alongside the number of high-confidence HHV-6B UMIs from the quantification pipeline. The number of HHV-6+ cells are barcodes annotated as cells before quality-control filtering in the original publication with at least one HHV-6 UMI. We note that many super-expressors contain high levels of mitochondrial and ribosomal transcripts, potentially due to these cells nearing apoptosis caused by viral infection/ reactivation.

For the ~1,000 donor PBMC dataset^37^, we similarly downloaded .fastq files from GEO and quantified with the resulting libraries for reads that pseudoaligned to the HHV-6B transcriptome. For either dataset, we did not detect a single HHV-6B read that was annotated by a barcode that was a cell in the processed metadata for either study though we did observe 6 total HHV-6B reads from the PBMC cohort (all mapping to different gel emulsion bead barcodes).

For the public CAR-T cell datasets, all .fastq files were downloaded through either the EGA portal^2^ or reanalyzed using a Terra workspace.^36^ Each dataset was processed with the same HHV-6B kallisto | bustools workflow. We observed no cells annotated in the complete metadata in the pre-infusion only study^2^, resulting in an annotation of 0 HHV-6+ cells from this dataset. For the 7 days post infusion blood draw in cohort 2, we observed a total of 26 unique 10x barcodes with at least 1 HHV-6B UMI whereas only 7 of the 26 barcodes were annotated as cells in the original study.^36^ While the high UMI abundance for the Axi-R-15 donor has no ambiguous interpretation, we note that for the other two donors (Tisa-R-32 and Axi-N-07) were pooled with the Axi-R-15 cells and then demultiplexed using genetic variants.^36^ As a consequence, we cannot exclude the possibility that the residual 1-2 UMIs detected in these 3 cells are from barcode contamination from the Axi-R-15 donor cells. Regardless, our characterization of the Axi-R-15 cells confirms the presence of HHV-6^hi^ cells in an FDA approved CAR-T cell product. For Axi-R-15, we note that the subject was not treated with steroids for ICANS and received one dose of dexamethasone for cytokine release syndrome on day +7 prior to the onset of ICANS. For the SJCAR19 analyses, the same kallisto| bustools pipeline was run locally at St. Jude to infer HHV-6 expression in the SJCAR19 cohort (cohort 3).

To confirm that HHV-6+ cells form the *in vivo* fusion product were from CAR-T cells, we assessed the presence of the CAR transgene via RNA (kallisto pseudo alignment to the axi-cell transcript) as well as noted host gene expression values (**Fig. 3; Extended Data Table 4**). The CAR ‘FACS’ cell was from the 10x channel that enriched for CAR+ cells via flow cytometry but due to the mechanisms of 10x Genomics library preparation protocol, the specific protein abundance for this individual cell is not discernable. The TCR α/β transcript number was the total number of UMIs aligning per barcode in the TCR vdj libraries.

### Expression of viral receptors across atlases

To evaluate other possible sources of HHV-6 reactivation and expression, we considered the landscape of cell types that express TNFRSF4 (OX40), the canonical receptor of HHV-6B, and CR2 (CD21), the canonical receptor of EBV. We downloaded processed count data and/or summary plots from the GTEx portal^44^, fetal development^45^, sorted resting and stimulated immune cells^46^, and endothelial cell lines^47^.

Analyses of these atlases revealed that OX40 is expressed specifically in activated CD4+ and CD8+ T cells in the immune system but had detectable levels across all tissues (**Extended Data Fig. 1**). When examining single-cell and single-nucleus expression across the GTEx atlas, we surprisingly observed consistent expression in endothelial cells across various tissues, including in the pancreas and brain. We further corroborated 15 of the top 25 cell types / tissues were endothelial. While the overall expression of OX40 appears to be 1-2 orders of magnitude higher in activated T cells, we suggest that the application of our atlas-evel analyses of the HHV-6B viral receptor as a possibility indicates that endothelial cells may be targets of active HHV-6B infection, either from the primary infection or from cell therapy-mediated reactivation. Moreover, we speculate that the brain endothelial cells expression OX40 mediates a possible mechanism underlying the encephalitis observed in severe infections/reactivation.

## Acknowledgements

We thank members of the Satpathy lab for helpful discussions. We are grateful to M. Green and G. Syal for assistance with public sequencing data. C.A.L. is supported by a Stanford Science Fellowship, a Parker Institute for Cancer Immunotherapy Scholarship, and a seed award from the Center for Human Systems Immunology at Stanford. This work was supported by a sponsored research agreement with Allogene Therapeutics. A.T.S. is supported by the Burroughs Wellcome Fund Career Award for Medical Scientists, the Parker Institute for Cancer Immunotherapy, a Pew-Stewart Scholars for Cancer Research Award, a Cancer Research Institute Lloyd J. Old STAR Award, and a Baxter Foundation Faculty Scholar Award. Part of the analysis was performed by the Center for Translational Immunology and Immunotherapy (CeTI^2^), which is supported by St. Jude Children’s Research Hospital.

## Author contributions

C.A.L. and A.T.S. conceived and designed the study with input from T.P., H.D., and R.G.M. C.A.L. lead all analyses and is responsible for the accuracy and reproducibility of the manuscript. Y.Y., K.M., K.D.S. performed scRNA-seq experiments and interpreted data. J.C.C., J.C.G., N.J.H., J.M.V., V.L., and A.K. supported the informatics analyses. K.M., J.C.C., L.P., G.G., M.V.M., A.C.T., P.G.T., S.G., and C.J.W. provided data and/or insights related to the in vivo HHV6 expression. G.Y., J.P., R.S., W.L., and A.S. designed, executed, and analyzed data related to HHV-6 in *in vitro* CAR-T data. A.M., R.A., T.P, and H.D. provided technical review, strategy, and guidance for *in vitro* CAR experiments. A.M.S., T.L.R., M.J.K., and R.G.M. interpreted experiments and guided clinical interpretation. C.A.L. and A.T.S. wrote the manuscript with input from all authors.

## Code and Data Availability

Raw and processed sequencing data used in this study is available at GEO Accession **GSE210063** with reviewer access token **kzmzgqsidfynlkd**. Code to reproduce all custom analyses in this manuscript is available online at https://github.com/caleblareau/hhv6-reactivation.

## Supplemental Materials

**Extended Data Table 1.** Top 5 BioSamples per virus from the Serratus screen.

**Extended Data Table 2.** All BioSamples mapping to a T cell annotation from the Serratus screen.

**Extended Data Table 3.** Complete list of host gene-HHV-6B expression correlation values.

**Extended Data Table 4.** Summary of UMI counts per feature in HHV-6 reactivated cells *in vivo*.

**Extended Data Table 5.** All BioSamples mapping to iPSCs from the Serratus screen.

**Extended Data Figure 1.**
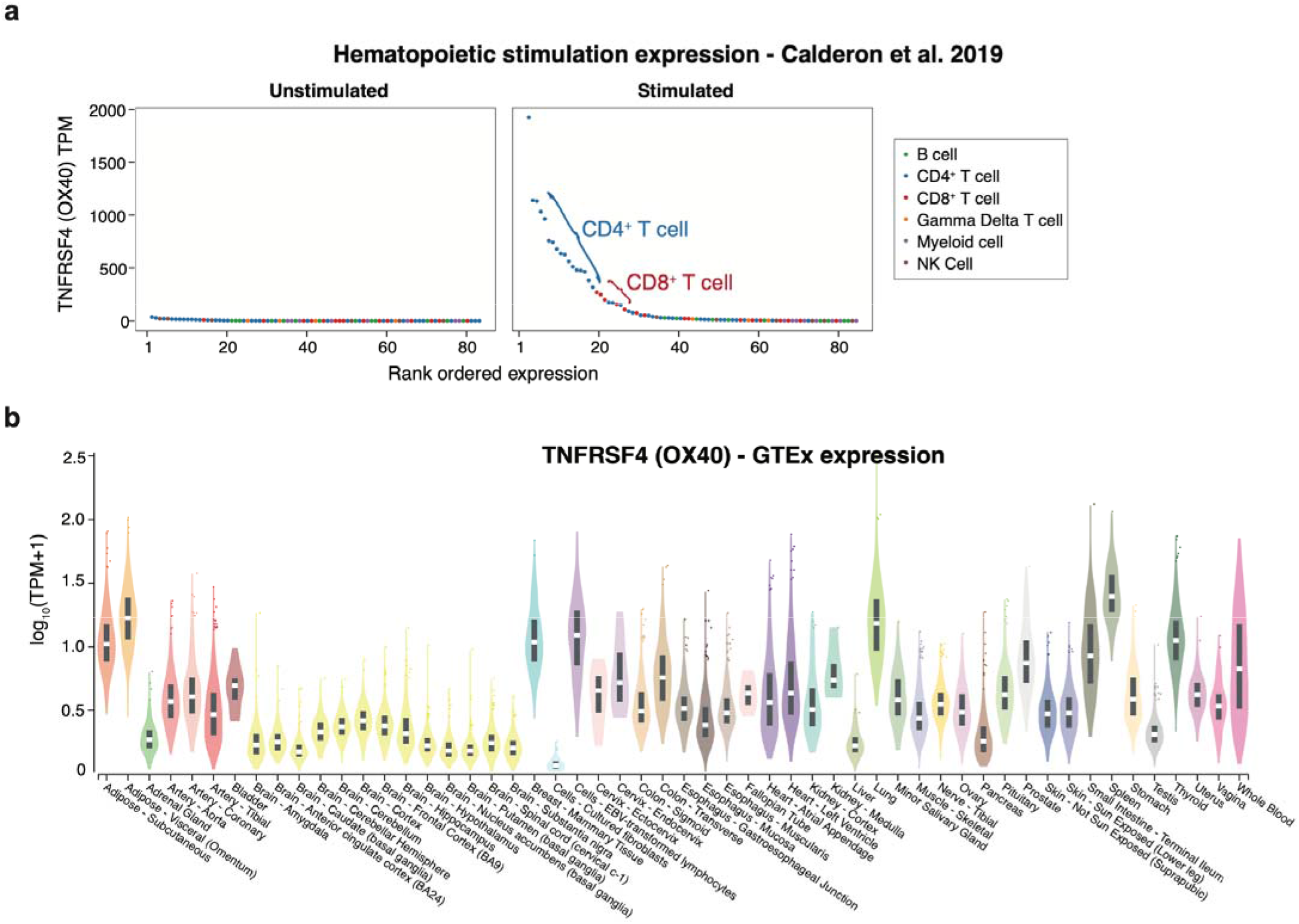
Characterization of OX40 expression in bulk sequencing experiments. **(a)** Expression of TNFRSF4 (OX40), the canonical receptor of HHV-6B, in unstimulated and stimulated immune cell populations.^46^ OX40 is not expressed in unstimulated immune cells but highly expressed in CD4 and CD8 T cells after activation/stimulation of CD3/CD28 and IL-2. **(b)** Broad expression of OX40 across healthy tissues from the GTEx bulk atlas.

**Extended Data Figure 2.**
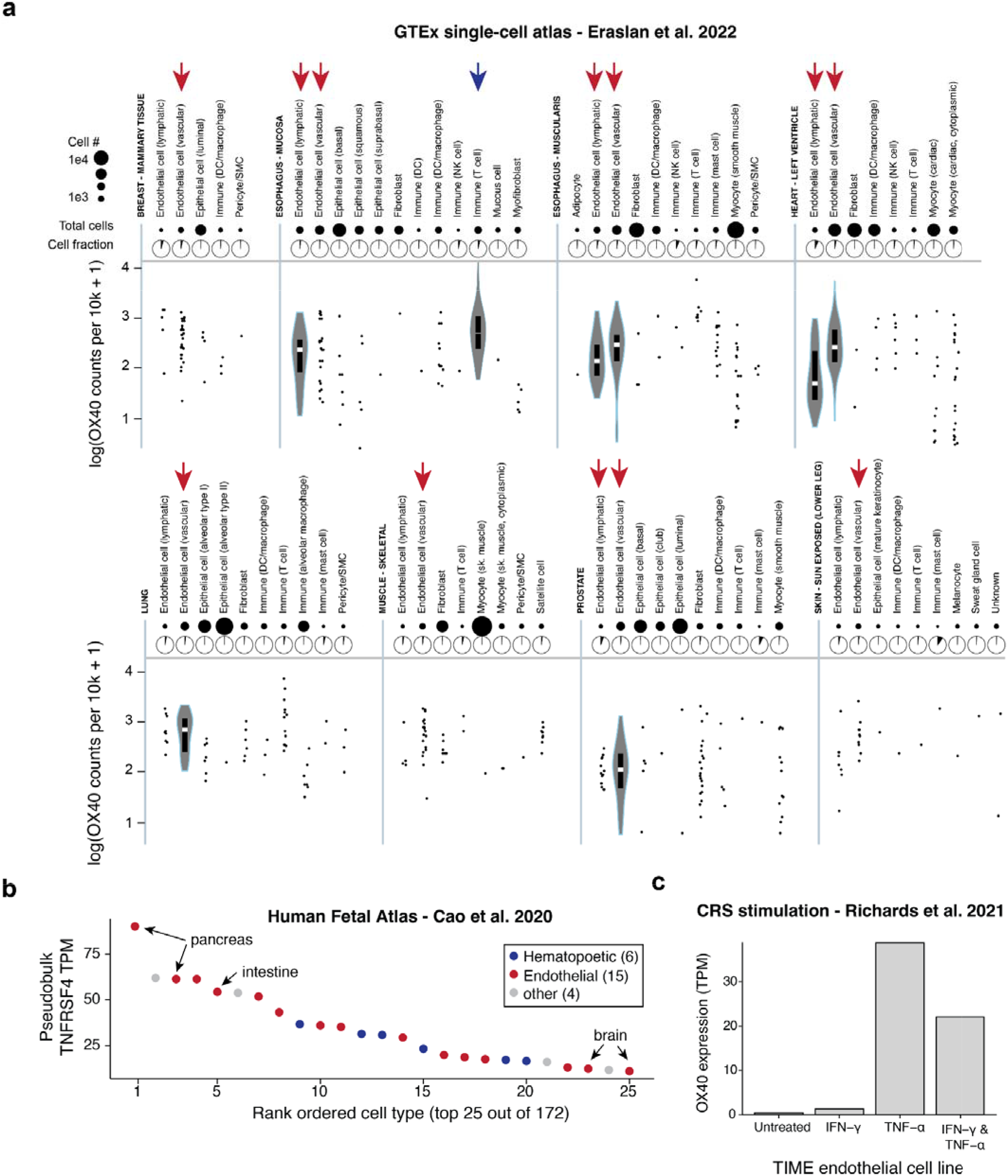
Characterization of OX40 expression in resting and stimulated endothelial cells. **(a)**Refinement of HHV-6B expression using the single-cell GTEx atlas^44^. **(b)** Pseudobulk expression of OX40 from the Human Fetal Atlas.^45^ **(c)** Induction of OX40 expression on endothelial cell lines in the presence of TNF-α; RNA-seq dataset from Richards *et al*.^47^

**Extended Data Figure S3.**
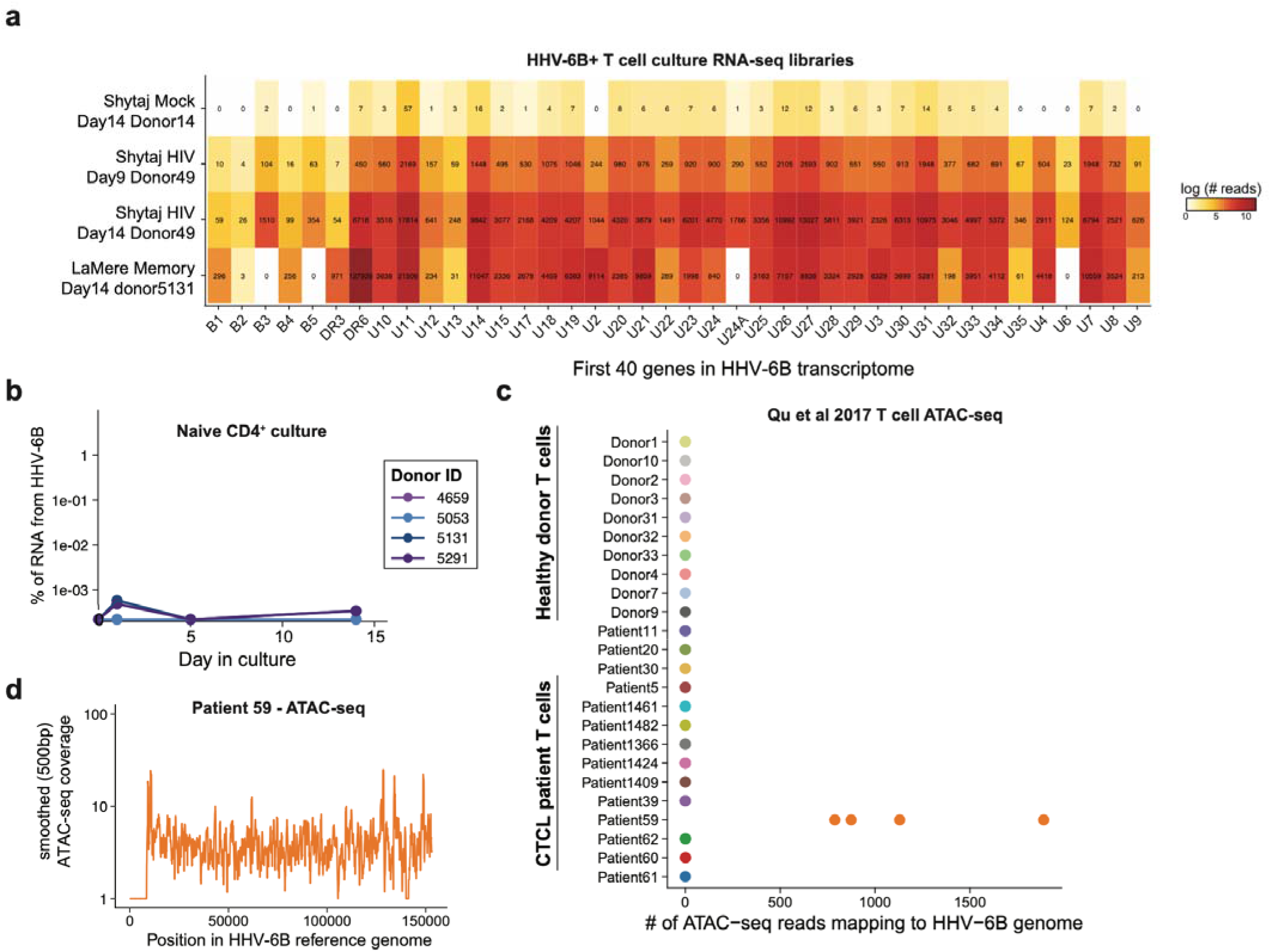
Supporting analyses for HHV-6 reactivation using Serratus. **(a)** Heatmap of HHV-6B transcripts across the four highest RNA-seq libraries from Serratus. Shown are the first 40 genes (based on genomic coordinate order) from the HHV-6B transcriptome and the number of reads that pseudoalign to each transcript. **(b)** Summary of naive CD4+ culture in the LaMere *et al*. dataset; compare to **Fig. 1e**. **(c)** Summary of HHV-6B expression in a previously reported ATAC-seq atlas,^20^ showing sorted T cells from Patient 59, an individual with CTCL, had detectable levels of HHV-6B DNA within cells. **(d)** Smoothed coverage (rollmean of 500 base pairs) over the four libraries from Patient 59, indicating coverage across the HHV-6B reference genome.

**Extended Data Figure 4.**
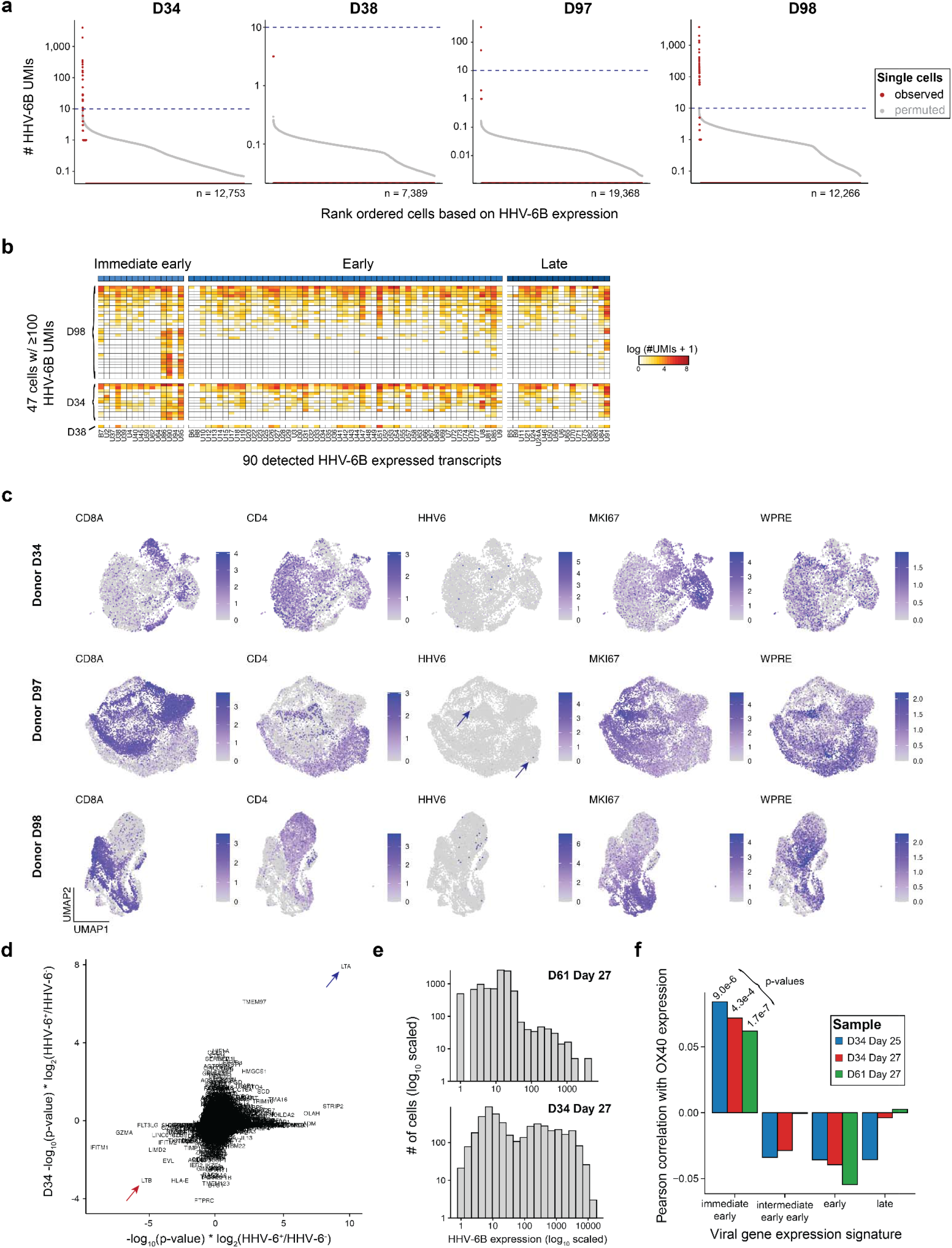
Supporting analyses for HHV-6 expression during *in vitro* CAR-T cell culture. **(a)** Summary of observed (red) and permuted (gray) HHV-6B expression for four donors at day 19 in culture. Dotted line is 10 UMIs, the threshold for a super-expressor. **(b)** Heatmap of HHV-6B expression for selected cells across 3 donors with detectable super-expressors. Columns are grouped based on HHV-6B gene programs (immediate early; early; late). **(c)** Uniform manifold approximation and projection for 3 samples, noting marker genes and HHV-6B UMI expression (log transformed). The WPRE feature indicates the presence of the CAR transgene. **(d)** Summary of HHV-6B +/-cells from differential testing for host factors. Arrows indicate lymphotoxin α (LTα/LTA) and downregulation of lymphotoxin β(LTβ/LTB). **(e)** Summary of HHV-6B expression in re-cultured samples for donors D61 and D34. **(f)** Correlation statistics of HHV-6B transcript signatures with OX40 expression across 3 recultured samples, including p-values from Pearson’s product moment correlation coefficient. The consistently positive, significant correlation statistic represents an association uniquely between OX40 expression and the immediate early HHV-6B gene signature.

